# Habitat preference of an herbivore shapes the habitat distribution of its host plant

**DOI:** 10.1101/156240

**Authors:** Nicolas M. Alexandre, Parris T. Humphrey, Andrew D. Gloss, Jimmy Lee, Joseph Frazier, Henry A. Affeldt, Noah K. Whiteman

## Abstract

Plant distributions can be limited by habitat-biased herbivory, but the proximate causes of such biases are rarely known. Distinguishing plant-centric from herbivore-centric mechanisms driving differential herbivory between habitats is difficult without experimental manipulation of both plants and herbivores. Here we tested alternative hypotheses driving habitat-biased herbivory in bittercress (*Cardamine cordifolia*), which is more abundant under shade of shrubs and trees (shade) than in nearby meadows (sun) where herbivory is intense from the specialist fly *Scaptomyza nigrita*. This system has served as a textbook example of habitat-biased herbivory driving a plant’s distribution across an ecotone, but the proximate mechanisms underlying differential herbivory are still unclear. First, we found that higher *S. nigrita* herbivory in sun habitats contrasts sharply with their preference to attack plants from shade habitats in laboratory choice experiments. Second, *S. nigrita* strongly preferred leaves in simulated sun over simulated shade habitats, regardless of plant source habitat. Thus, herbivore preference for brighter, warmer habitats overrides their preference for more palatable shade plants. This promotes the sun-biased herbivore pressure that drives the distribution of bittercress into shade habitats.

## Introduction

Abiotic gradients shape fine-scale patterns of plant distributions across the landscape (Whittaker 1967), but consumers, such as insect herbivores, can play a major role as well (Harley 2003; Fine et al. 2004; Maron and Crone 2006). Herbivores can drive plant distributions by reducing plant fitness in some habitats more than others, and this can occur both because of (A) differential impacts of a given level of herbivory, or (B) differential rates of herbivory itself. Abiotic variation can impact susceptibility to herbivores (e.g. via growth–defense trade-offs; Fine et al. 2004, 2013), which leads herbivores to promote plant habitat specialization. But when the total intensity of herbivory itself also varies across habitats (e.g. Louda and Rodman 1996, Fine et al. 2006, 2013), it can be difficult to discern whether plant-centric or herbivore-centric mechanisms are responsible. Distinguishing among the mechanisms that shape herbivore distributions is vital for understanding their impacts on plant distributions as well as the likely responses of both herbivores and plants to changes to their abiotic environments.

What distinguishes plant-from herbivore-centric mechanisms is whether the plant or the herbivore response to abiotic gradients takes precedence in shaping realized patterns of herbivore pressure across habitats. Differential herbivore pressure between habitats can arise because herbivores seek out higher quality hosts, and in this case, plant-centric mechanisms (e.g. reduced plant defenses) ultimately shape herbivore distributions. Thus, the habitat effect on the herbivore is indirect, mediated instead by habitat-specific variation in plant traits. In contrast, herbivore-centric hypotheses posit that herbivores are more abundant in, or seek out, favorable abiotic habitat conditions (Huffaker and Kennett 1959), independent of how plant traits vary across habitats. In this case, herbivore habitat tolerance and/or preference is directly shaped by abiotic conditions, which creates enemy-free space exploitable by plants. Here we addressed how plant-versus herbivore-centric factors impact the habitat-specific herbivory pressure that is responsible for shaping the habitat distribution of a native subalpine plant.

Studies on bittercress (Brassicaceae: *Cardamine cordifolia*) in the Rocky Mountains of North America were among the first to explore how fine-scale variation in herbivory shapes plant fitness and abundance across habitats (Collinge and Louda 1988, 1990; Louda 1984; Louda and Rodman 1983, 1996), and this system is a textbook example of an herbivore-limited plant distribution (Ricklefs and Miller 2000). *Scaptomyza nigrita* flies (Drosophilidae) are a major herbivore of bittercress: female adults make feeding punctures (‘stipples’) and oviposit in leaves, and the leaf-mining larvae can defoliate up to 70% of leaf area in sun habitats (Collinge and Louda 1988). Herbivory is higher in sun habitat, and the fitness effects are strong enough to drive bittercress into the shade (Louda and Rodman 1996). Surprisingly, the proximate drivers of differential herbivore pressure across this ecotone remain largely unknown. Collinge and Louda (1989) found that plants growing in sun habitats suffer higher herbivory in part because of plant phenology: immediately after snowmelt, flies only have access to plants that have emerged in sun habitats. However, flies are still abundant weeks after this period, and plants in the shade still suffer low herbivory (Collinge 1989). We sought to address why this is the case.

Both plant- and herbivore-centric mechanisms can be proposed to explain sun habitat-biased herbivory (*sensu* Louda and Rodman 1996). In addition to their earlier availability, plants in sun habitats may be less resistant, and thus more attractive or palatable to *S. nigrita*, than those inshade habitats. Under this plant-centric hypothesis, higher plant quality in the sun would cause sun-biased herbivory. Several lines of evidence are consistent with this hypothesis: bittercress from sun habitats can have lower glucosinolate (GSL) content, the precursors of toxic mustard oils (isothiocyanates) (Louda and Rodman 1996), and GSL-enriched bittercress can deter adult female *S. nigrita* and harm their larvae (Humphrey et al. 2016). Additionally, foraging adult females are more active and abundant in sun versus shade habitats (Louda and Rodman 1996), which could arise if *S. nigrita* seek out higher quality plants as they forage across the ecotone.

Under an alternate hypothesis, sun habitat-biased herbivore abundance—and thus overall herbivore pressure—could arise because *S. nigrita* are attracted, or restricted, to sun habitats due to abiotic habitat features. Under this herbivore-centric hypothesis, higher herbivore pressure on sun plants arises because there are simply more flies in sun. Thus, a direct effect of the abiotic environment on herbivore behavior would release shade-associated bittercress from herbivore pressure, and this mechanism can operate with or without reinforcement from plant phenotypes. Whether the proximate driver of variation in herbivore pressure across this ecotone arises from plant- or herbivore-centric mechanisms has important implications for the types of natural selection faced in each habitat by both plants and herbivores (Fine et al. 2006).

Here we provide a test of these alternate proximate causes of sun-biased herbivory in bittercress. We first revisited whether *S. nigrita* herbivore pressure is higher in the sun relative to shade habitats by conducting field herbivory surveys. Second, we tested whether *S. nigrita* preferentially forage on shade- or sun-source plants by offering *S. nigrita* females a choice of the two bittercress types under laboratory conditions. We then tested the hypothesis that abiotic features of sun and shade habitats drive feeding and oviposition behavior by manipulating light and temperature in a series of choice trials conducted under laboratory and field settings, using plants from both sun and shade habitats. Altogether, our experiments support an herbivore-centric behavioral explanation for the sun-biased herbivory pattern that shapes the habitat distribution of this textbook native interaction system.

## Materials and Methods

### Herbivory surveys

All experiments were conducted between 2010 and 2015 at the Rocky Mountain Biological Laboratory (RMBL) in Gothic, CO, USA. In 2011, we conducted field surveys of herbivore damage on bittercress in 8 sun habitats (open meadows) and 7 shade habitats (under dense evergreen tree canopies; Appendix S1: Fig. S1, Table S1). We recorded adult *S. nigrita* feeding punctures (stipples), larval mines, and leaf area of 2 basal leaves from each of 10 ramets from all 15 bittercress patches.

We modeled feeding punctures made by adult females (stipples) and larval mine counts using zero-inflated (ZI) negative binomial (ZINB) generalized linear mixed models. Zero-inflation (i.e. under-dispersion) describes a notable excess of observed zero counts relative to the expected zero counts arising under non-truncated Poisson or NB processes (Zuur at al. 2009). In biological terms, zero-inflation can arise from patchily distributed herbivores and clustered feeding behavior, resulting in many un-damaged leaves (zero counts) even while plants that are damaged may tend to have large amounts of damage (a typical negative binomially distributed pattern for parasites). Thus, herbivory intensity data can simultaneously exhibit both under- and over-dispersion, and this can generate lack-of-fit and biased parameter estimates if ignored. Such was the case with our herbivore survey data (Appendix 3: Fig. S1).

We therefore constructed models that explicitly modeled ZI, as a single parameter and as a function of habitat type, using the canonical logit link function in a binomial GLM. The non-ZI count class was simultaneously estimated via NB GLM with a log link function, with habitat (sun vs. shade) and leaf area (mm^2^) included as population-level (fixed) terms, and site ID and plant ID as group-level (random) intercept terms. Coefficients were estimated via maximum likelihood using R v3.3.3 (R Core Team 2017) package *glmmTMB* v. 0.2.0 (Brooks et al. 2017, Magnusson et al. 2017).

In addition, the flexibility of package *glmmTMB* allowedustoestimatetheNBdispersionparameter (*ϕ*).as a function of source type, with and without simultaneous accounting for ZI. Both parameters help account for unequal variance across residuals and improve goodness-of-fit (higher values of *ϕ*.correspond to *lower* variance in the present NB variance formulation; Appendix 3). We evaluated fixed effects by step-wise model reduction (Zuur et al. 2009) and model comparisons via Akaike’s Information Criterion (*AICc*) (Hurvich and Tsai 1989). We compared best-fitting models to those with additional parameters that estimated ZI and NB dispersion as a function of site type using *AICc* and with predictive model checks to inspect goodness-of-fit. Model details, diagnostics, and comparisons can be found in Appendix S3.

### Host choice experiment I: Sun versus shade-derived bittercress

In 2010 we tested whether *S. nigrita* adult females prefer feeding on individual bittercress derived from sun or shade habitats. We transplanted bolting bittercress plants from the field into soil within plastic pots in the laboratory under fluorescent lighting (16:8 light:dark) for < 24 h. In each of 8 replicates, we randomly assigned 2 shade-derived and two sun-derived bittercress plants to the 4 corners of a mesh 35.5 x 35.5 x 61 cm cage (<u>livemonarch.com</u>) (see Appendix S2: Fig. S1A). All leaves were un-mined, and we subtracted pre-existing stipple damage from final counts. Four field-collected adult female flies were introduced into each cage and allowed to feed for 24 h, after which stipples and eggs were counted with a dissecting microscope.

To control for differences in plant architecture between sun- and shade-derived bittercress, we conducted a detached leaf assay using cauline leaves clipped from the flowering stalk of first or second lowest position of plants from sun or shade habitats. For each of 15 replicate trials, 2 leaves each from sun and shade plants were inserted by their petioles into a half liter-sized transparent plastic container filled to a depth of 1.5 cm with 2% Phytoblend (Caisson Laboratories, Logan, UT). Leaves were randomly assigned to positions for each assay container, which was closed with a plastic and mesh lid (see Appendix S2: Fig. S1B for a schematic). We introduced one field-caught adult female fly into each container and allowed it to forage for 24 h, after which we counted stipples and eggs. No flies were used for multiple trials.

For both assays we modeled stipple and egg counts with NB mixed models using *glmmTMB*, with plant habitat (sun vs. shade), number of cauline leaves (for whole-plants), and leaf width(for detached leaves; mm) as fixed effects, and cage ID (i.e. replicate assay) as a group-level random intercept term. We evaluated support for habitat-specific estimates of*θ*asabove.We assumed that adult females *S. nigrita* flies had sufficient time to potentially visit all available plant tissue; thus, *a priori* we favored models without ZI terms.

### Host choice experiment II: Effects of light and temperature

In 2014 and 2015 we conducted choice experiments in the laboratory and field to decouple the effects of light and temperature on *S. nigrita* foraging behavior. At two-day intervals, we conducted 6 trials using both sun-warmed and shade-cooled large mesh cages (35.5 x 35.5 x 185 cm) where one side of each cage was randomized to receive a lighting treatment. Ten undamaged bolting bittercress were collected near RMBL along the Copper Creek Drainage (CCD) (Appendix S1: Fig. S1) and maintained in the laboratory for ≤ 4 d prior to each trial. Four leaves from each of the 10 plants were detached at the petiole and randomized to each of 4 experimental conditions (2 cage-level temperature treatments 2lightenvironmentspercage).Leaf petioles were fixed with a moist paper towel in 100 mm petri dishes placed at either end of the cages (Appendix S2: Fig S2). Ten *S. nigrita* adult females were collected along the CCD (Appendix S1: Fig. S1), released into the middle of each cage, and allowed to forage for 24 hours starting at 1100 h. Additional details on the methods and design of these cage experiments can be found in Appendix S2.

For the 2014 laboratory choice trials, 2 large mesh cages were placed into temperature-controlled environmental chambers that were either cooled or held at ambient temperature (∽16°C and∽21°C, respectively; Appendix S4: Fig. S1). Plants, leaves, and flies were collected and utilized as above, and flies were allowed to feed for 8 hours (1100–1900 h) during each trial. We carried out similar trials in 2015 but in a single environmental chamber at 2 day intervals, alternating between approximately 20 °C and 24 °C (Appendix S4: Fig. S1). Leaves were obtained along the CCD (Appendix S1: Fig. S1) and were randomized across treatments as before. Baseline temperatures in 2015 were elevated by 4 °C relative to 2014 (Appendix S4: Fig. S2). In addition to stipples, we counted eggs deposited by *S. nigrita*, which were not counted in 2014 because our experiment began later in the season when adult females were not often gravid.

We modeled stipple and egg counts using NB mixed models using *glmmTMB* with the following fixed effects: leaf width (mm^2^), leaf position along stem from which it was removed (‘position’), light environment (light vs. dark,), temperature (warm vs. cool,), and an interaction term between temperature and light environment. We modeled between-trial, between-room, between-cage, and between-side-of-cage effects as a series of nested random intercept terms. For 2015 trials, we also included plant source habitat (sun vs. shade) as a fixed effect. Finally, we evaluated support for inclusion of condition-specific NB dispersion parameters via *AICc* comparisons to the best-fit model without such terms, as above.

For all analyses, statistical significance of fixed effects was assessed at the *p*<0.05 level via asymptotic Wald’s *z* tests. Average differences in herbivore damage counts reported in the results are predicted mean (with predicted 95% confidence interval) of the response variable generated via 1000 simulations from the best-fitting model, using the maximum likelihood point estimate of model coefficients.

## Results

### Herbivory surveys

Across 15 field sites, naturally-occurring stippling and leaf miner damage from *S. nigrita* was strongly biased towards bittercress in sun habitats compared to shade habitats (Fig. 1A). Compared to plants in shade habitats, leaves of plants in the sun showed an overall 5-fold higher stipple abundance (16.1 [12.5–20.4, 95% CI] versus 3.2 [2.3–4.3] mean stipples per leaf, *p*<0.001; Table 1). We estimated that 29% [20–37%, 95% CI] of leaves in the shade avoided stippling altogether, compared to only 9% [5–14%] of leaves in the sun. Sun plants also had a>40-fold higher overall leaf mine abundance (3.7 [2.0–6.7] vs. 0.09 [0.03–0.21] mean mines per leaf; *p*<0.001; Table 1), and 93% [88–98%] of leaves in shade habitats had no leaf mines at all, compared to only 29% [19–40%] in the sun.

**Fig. 1.**
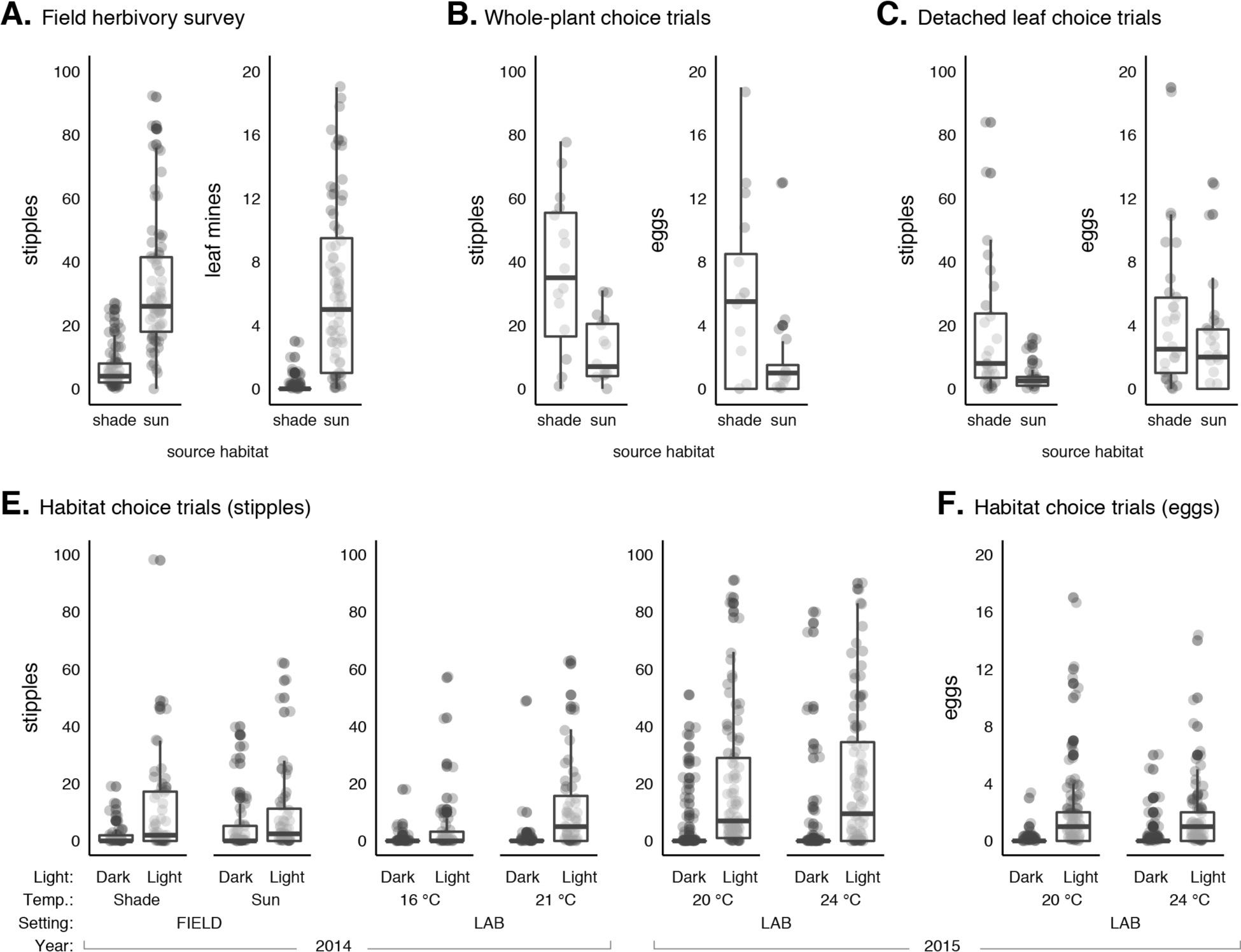
Choice experiments reveal that habitat selection, and not host selection, by foraging *Scaptomyza nigrita* underlies the pattern of sun-skewed herbivory in nature, despite higher palatability of bittercress from the shade. **(A–B)** Herbivory is higher on bittercress in sun versus shade habitats, but female *S. nigrita* prefer shade-grown over sun-grown bittercress when given a choice under uniform lighting. (**A**) Herbivory field surveys show higher stipples and mines on bittercress in sun vs. shade habitats. (**B**) Adult female *S. nigrita* stippled and laid more eggs in bittercress derived from shade versus sun habitats in laboratory choice trials. Statistical results are presented in Table 1. (**C–D**) Female *S. nigrita* stippled more (**C**) and laid more eggs (**D**) in bittercress leaves in simulated sun compared to shade habitats in field and laboratory choice trials. Choice trials between light and dark sides of assay cages were conducted at two temperatures (see Appendix S4: Fig. S1 for full temperature profiles), indicated below sub-plots. Eggs were counted only for trials in 2015 (see Materials and Methods). Statistical results are presented in Table 1.

**Table 1.**
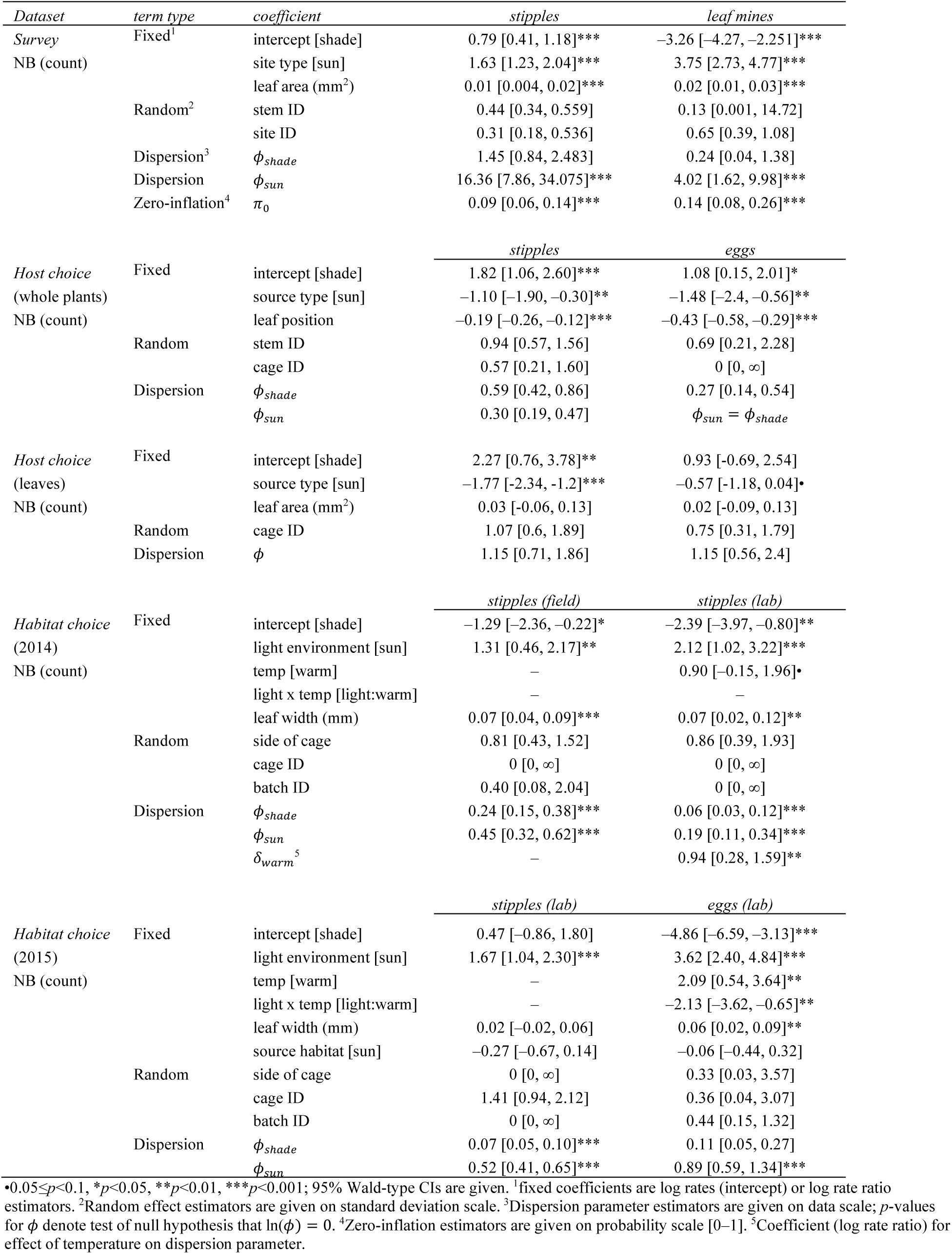
Coefficient estimates for herbivory models across habitats and choice trials.

### Host choice experiment I: Sun versus shade-derived bittercress

In contrast to patterns revealed in the herbivory survey, *S. nigrita* female flies strongly preferred feeding and laying eggs on bittercress from shade habitats when given a choice of whole plants from shade or sun habitats under uniform light conditions (Fig. 1B). Compared to shade-source plants, overall stipple abundance on sun-source plants was 60% lower (1.4 [0.6–3.1] versus 4.5 [1.9–9.9] mean stipples per leaf, *p*<0.01). Female flies left sun-source leaves free of stipples 71% [61–80%] of the time, compared to only 43% [30–55%] of the time for shade-source plants. Overall egg abundance was 75% lower on sun-source plants (0.13 [0.05–0.25] versus 0.64 [0.32– 1.10] mean eggs per leaf, *p*<0.01; Table 1), and 93% [88–96%] of sun-source leaves remained free of eggs compared to 80% [74–87%] of shade-source leaves.

When individual leaves were offered instead of whole plants (Fig. 1C), overall stipple abundance was >80% lower for sun-source leaves (4.8 [1.8–10.9] versus 27.9 [12.1–60.8] mean stipples per leaf, *p*<0.001). While a similar proportion of leaves from each habitat received zero eggs (shade: 24% [10–40%], sun: 35% [20–53%]), overall egg abundance for sun-source plants was 45% lower than for shade-source plants (2.5 [1.2–4.3] versus 4.4 [2.3–8.0] mean eggs per leaf). This difference was marginally significant (*p*=0.065; Table 1), and an intercept-only model was favored via model comparisons (△*AIC*c= –1.2).

### Host choice experiment II: Effects of light and temperature

In the 2014 field trials, *S. nigrita* strongly preferred feeing in the lighted sides of the cages over the unlighted sides of the cages (Fig. 1E). Overall stipple abundance was 4-fold higher on plants under lights compared to those under shade (18.1 [6.3–52.8] versus 4.0 [1.5–9.9] mean stipples per leaf, *p*<0.001; Table 1). We detected no effect of cage warming, and removal of temperature terms was statistically favored via model comparisons (△*AIC*c = –1.2). *S. nigrita* preference for feeding in the light was strong in both 2014 and 2015 laboratory choice trials (Fig. 1E): average stipple abundance was 8-fold higher in 2014, and 5-fold higher in 2015, on leaves in the light compared to the unlighted sides of the cages (2014: 9.7 [4.5–19.3] versus 1.1 [0.3–2.9] mean stipples per leaf; 2015: 38.5 [13.8–93.9] versus 6.9 [1.5–20.7]; both *p*<0.001, Table 1). For 2014, warmed cages exhibited a marginally significant 2.5-fold increase in stipple abundance (*p*=0.094, Table 1), while for 2015 trials we detected no temperature effect and removal of this term was statistically favored (△*AIC*c= 2.0).When both years’datawerepooled,temperatureremained non-significant, and all other results were qualitatively unchanged (Appendix S3: Table S2).

In warmed cages, egg abundance was 4.5-fold higher on leaves under lights than those in the unlighted sides of cages (Fig. 1F; 1.6 [0.8–2.7] versus 0.36 [0.1–0.8] mean eggs per leaf, respectively; *p*<0.001, Table 1). Cooler temperatures reduced egg laying 6-fold in the dark (down to 0.05 [0–0.12] mean eggs per leaf) but had no effect in the light (light-by-temperature interaction term *p*<0.01; Table 1). Notably, plant source habitat (sun vs. shade) did not impact stippling (*p*>0.2) or egg (*p*>0.7) abundances (Appendix 5 Fig. S1), and inclusion of this term was never supported via model comparisons using *AICc*.

## Discussion

Here, we report evidence of a proximate explanation for a textbook case of an herbivore-driven habitat distribution: insect behavioral taxis strongly biased herbivore foraging towards sun habitats, causing the increased herbivore pressure on sun plants that drives bittercress into the shade. Consistent with the pattern found by Louda and Rodman (1996), we found that overall herbivore pressure was higher on bittercress naturally growing in sun than in shade habitats (Fig. 1A). Contrary to this sun-biased herbivory pattern in the field, *S. nigrita* strongly preferred bittercress from shade when given a choice (Fig. 1B–C). But when we manipulated the abiotic conditions under which herbivores foraged, a strong preference for bright habitats emerged (Fig. 1E) that overrode their preference for plants from shade habitats. In fact, warmer temperatures and high light levels combined to drive herbivory into simulated sun habitats: the fewest eggs were laid on leaves away from lights and in cooler cages (Fig. 1F). Thus, the distribution of herbivore pressure across the sun/shade ecotone that drives bittercress into the shade (Louda and Rodman 1996) results from a direct effect of the abiotic environment on herbivore behavior.

A variety of adaptive and non-adaptive explanations can be posited to explain a strong habitat preference for feeding and oviposition by *S. nigrita*. In particular, oviposition preference should reflect the habitat distribution where larvae have the greatest probability of survival (Craig et al. 2000). While adult herbivore feeding and oviposition preference does not necessarily predict larval performance (Craig et al. 1999; Craig et al. 2000), evidence from a recent study supports the notion that shade-source bittercress are, in fact, *higher* quality for *S. nigrita* than sun-source plants: short-term larval performance was higher in shade-source plants compared to sun-source plants when both were re-grown in shade habitats (Humphrey et al. 2018). This is consistent with preferences of adult *S. nigrita* for shade-derived plants reported in this study and adds to the evidence that *S. nigrita* foraging preferences for certain plant tissues over others generally reflect differences in plant quality for larvae (Humphrey 2016, 2017). Measuring fitness through the entire life cycle will be essential for testing the hypothesis that choosing shade-source plants is (or would be) adaptive for *S. nigrita*.

Alternatively, a non-adaptive (or mal-adaptive) explanation is that herbivore attraction to light is too strong to permit foraging on the higher quality plants in nearby shade habitats. We regard the constraint hypothesis as intriguing but implausible, because phototactic behavior can vary plastically and genetically both between and within species of drosophilids (Gorostiza et al. 2016). The fact that it persists in *S. nigrita* suggests that there may be benefits to feeding in warm, sunny habitats at >3000 m. in elevation that outweigh any advantages to feeding on the more palatable host plants in the shade. Cool temperatures restrict the ability of insects to oviposit on available host plants, even when abundant, because temperatures are too low for flight (Kingsolver 1989). This may explain why insects are often restricted to sunny habitats (Huffaker and Kennett 1959), areas experiencing sunny weather (Whitman 1987), or areas within a plant exposed to the sun, regardless of plant quality (Casey 1993). Separate from thermal tolerance, perception may simply be more efficient in the sun, where flies may rely on visual cues or the “clumped” distribution of host plants (Wallace 1958, Vernon and Gillespie 1990). Exploring how thermal tolerance, insect perception, and variation in host-plant quality interact to reinforce herbivore habitat preferences is a promising future research direction in this system.

Establishing whether herbivore-or plant-centric mechanisms shape the distribution and/or abundance of herbivores is crucial for understanding the nature of the selective forces that promote habitat specialization. Even in well-studied systems (e.g., Bruelheide and Schiedel 1999; Fine et al. 2004, 2006, 2013), the mechanisms (plant-centric or herbivore-centric) responsible for the differential herbivore pressure that shapes plant habitat distributions have been difficult to ascertain. In the Amazon, for example, insect herbivores promote habitat specialization by polarizing the allocation strategies best suited towards resource-rich clay versus resource-poor white-sand habitats (Fine et al. 2004). Clay habitats which favor increased plant growth over anti-herbivore defenses also tend to have higher herbivore pressure (Fine et al. 2006) and herbivore abundances (Fine 2013), but whether this is a consequence of reduced plant defenses or herbivore-centric mechanisms has not been addressed. Our study on a textbook system is novel because it provides a direct test of whether habitat-biased herbivory arises from herbivores tracking plant quality or from herbivore behaviors that arise independent of (or even in spite of) differences in plant quality. Our work suggests that the abiotic environment has a direct effect on maintaining enemy-free space in shade habitats. This, in combination with early-season plant phenological escape from herbivory (Collinge and Louda 1989), drives the distribution of bittercress towards shade habitats. As a consequence, strong herbivore habitat preference—regardless of its adaptive value for the insect—likely alters the nature and strength of selection on plant defense strategies across this sun/shade ecotone (Humphrey et al. 2018).

## Acknowledgments

We thank Ian Billick (RMBL), Jennifer Reithel (RMBL), Kailen Mooney (UC-Irvine), Carol Boggs (University of South Carolina), Mary Price (RMBL), Nick Waser (RMBL), Brian Enquist (University of Arizona), and Tom Miller (Rice University) for feedback. Financial support was from the National Science Foundation (NSF DEB-1256758 to NKW; DEB-XXXXXX to ADG); John Templeton Foundation (41855 to NWK); the National Institutes of Health (R35GM119816 to NKW); and the RMBL for research-(NKW), graduate-(PTH and ADG) and undergraduate-fellowships (HA and FJ).

## Appendix 1

### Characteristics of source locations for bittercress herbivory survey

At each site, we recorded leaf area of all sampled leaves, photosynthetically active radiation (PAR) using a light meter (Spectrum Technologies, Inc.), percent canopy cover using a densiometer, diameter at breast height (*dbh*) of the four largest trees within four meters, and latitude, longitude, and elevation using a GPS unit (Garmin) (Table S1). Environmental variables at each site were compared using one-way ANOVA. Sun habitats had higher average PAR and % open canopy than shade habitats (both *p* < 0.001) and did not systematically differ in elevation (*p* > 0.8, Table S2).

**Fig. S1.**
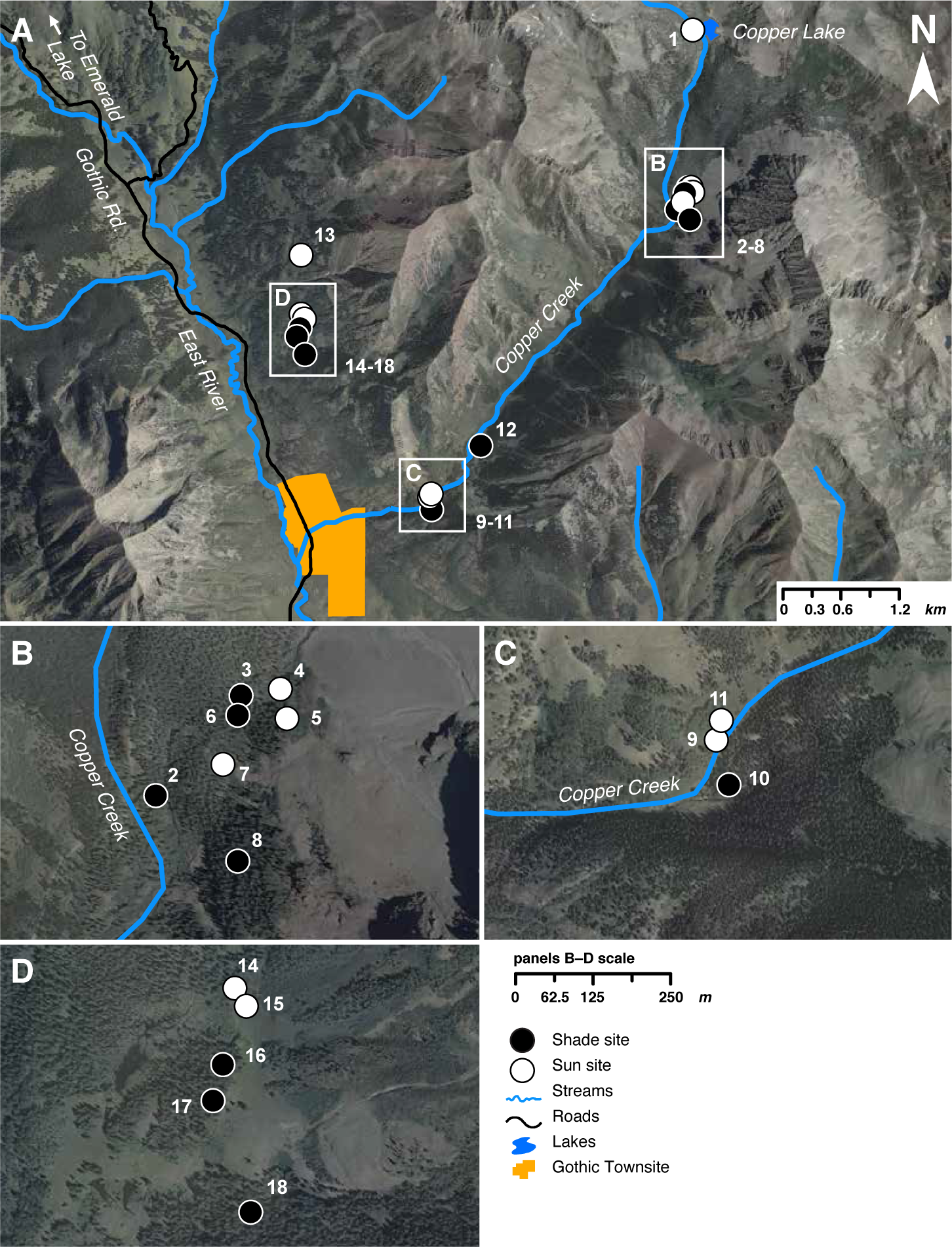
Map of source sites used in the herbivory surveys in the East River Valley and Copper Creek drainages, near the RMBL in Gothic, CO. **A**. Base map showing all sites within region (1:48,000). **B–D**. Maps showing detail of site locations (all same scale, 1:7500).

**Fig. S2.**
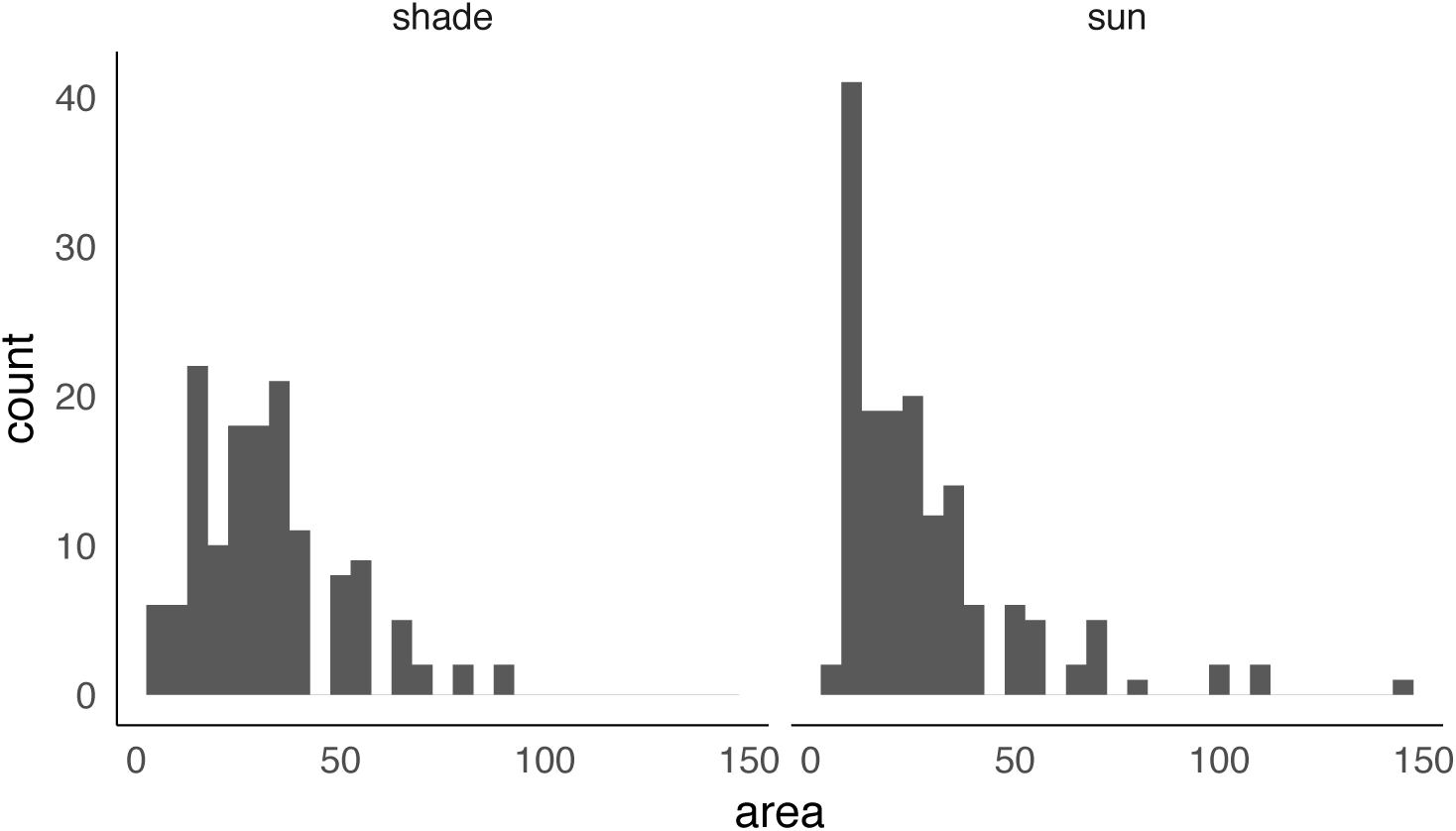
Distribution of bittercress leaf area (mm^2^) observed across n=298 leaves sampled across 8 shade and 7 sun habitats (Table S1).

**Table S1.** Attributes of sites (*n*=15) used for herbivory survey.

| Site # | Date sampled | Lat. | Long. | Elevation (m) | Soil Moisture | Light Environment | PAR ($\mu\text{mol}\cdot\text{s}^{-1}\cdot\text{m}^{-2}$)<sup>1</sup> | % Canopy Open | Average DBH (cm) |
| --- | --- | --- | --- | --- | --- | --- | --- | --- | --- |
| 4 | 11-Jul-11 | 38.9900829890862 | -106.943450029132 | 3253 | Very Wet | Sun | 2030 | 82.94 | 0 |
| 5 | 11-Jul-11 | 38.9896982273227 | -106.9432894133 | 3261 | Very Wet | Sun | 2034 | 95.42 | 0 |
| 6 | 11-Jul-11 | 38.9897805369006 | -106.94429611421 | 3244 | Moist Loamy | Shade | 130 | 4.16 | 83.25 |
| 7 | 11-Jul-11 | 38.9890385630266 | -106.944483622637 | 3220 | Very Wet | Sun | 2134 | 99.84 | 0 |
| 8 | 11-Jul-11 | 38.9876108211294 | -106.944190554921 | 3250 | Moist Loamy | Shade | 28 | 3.38 | 81.75 |
| 9 | 12-Jul-11 | 38.9607056556777 | -106.973679741745 | 3023 | Very Wet | Sun | 2026 | 88.14 | 0 |
| 10 | 12-Jul-11 | 38.9600423351315 | -106.973476684125 | 3013 | Wet Loamy | Shade | 35 | 5.72 | 72.25 |
| 11 | 12-Jul-11 | 38.9609507737634 | -106.973571138078 | 3994 | Very Wet | Sun | 1933 | 86.84 | 0 |
| 12 | 12-Jul-11 | 38.9655594715881 | -106.968516958823 | 3994 | Very Wet | Shade | 32 | 3.9 | 101 |
| 13 | 13-Jul-11 | 38.9828665210329 | -106.989822099201 | 4058 | Very Wet | Sun | 1780 | 96.2 | 0 |
| 14 | 13-Jul-11 | 38.9774145794955 | -106.98986568487 | 4082 | Very Wet | Sun | 1870 | 88.4 | 0 |
| 15 | 13-Jul-11 | 38.9771831850899 | -106.989697600334 | 4057 | Very Wet | Sun | 1850 | 99.84 | 0 |
| 16 | 13-Jul-11 | 38.976356744414 | -106.990078474138 | 4052 | Very Wet | Shade | 54 | 4.68 | 98 |
| 17 | 13-Jul-11 | 38.975757817399 | -106.990327194402 | 4046 | Moist Loamy | Shade | 60 | 4.16 | 140.5 |
| 18 | 13-Jul-11 | 38.974195183581 | -106.989498527449 | 4051 | Very Wet | Shade | 40 | 7.28 | 78.75 |

<sup>1</sup> Light meter was positioned above the center of each source collection plot, at times without cloud cover between 1:00 pm and 3:00 pm during the week of July 17, 2011.

**Table S2.** Environmental attributes of sun and shade sites for herbivory survey.

| *Site attributes* | Sun sites<br>(*n* = 8) |  | Shade sites<br>(*n* = 7) |  | ANOVA |  |
| --- | --- | --- | --- | --- | --- | --- |
| | $\mu$ | se | $\mu$ | se | *F* | *P* |
| PAR ($\mu\text{mol}\cdot\text{s}^{-1}\cdot\text{m}^{-2}$) | 1959.7 | 37.0 | 89.0 | 29.4 | 1564 | $<10^{-10}$ |
| % Canopy Open | 93.1 | 2.2 | 5.3 | 0.6 | 1527 | $<10^{-10}$ |
| DBH (*cm*) | 0.00 | 0.00 | 87.5 | 12.2 | 51.77 | $<10^{-10}$ |
| Elevation (*m*) | 3601 | 146 | 3568 | 150 | 0.025 | 0.875 |

| *Sample attributes* | Sun leaves<br>(*n* = 157) |  | Shade leaves<br>(*n* = 140) |  | ANOVA |  |
| --- | --- | --- | --- | --- | --- | --- |
| | $\mu$ | se | $\mu$ | se | *F* | *P* |
| Leaf size (*mm*) | 4.98 | 0.15 | 5.47 | 0.13 | 5.95 | 0.015 |

## Appendix 2

### Design schematics for choice experiments

**Fig. S1.**
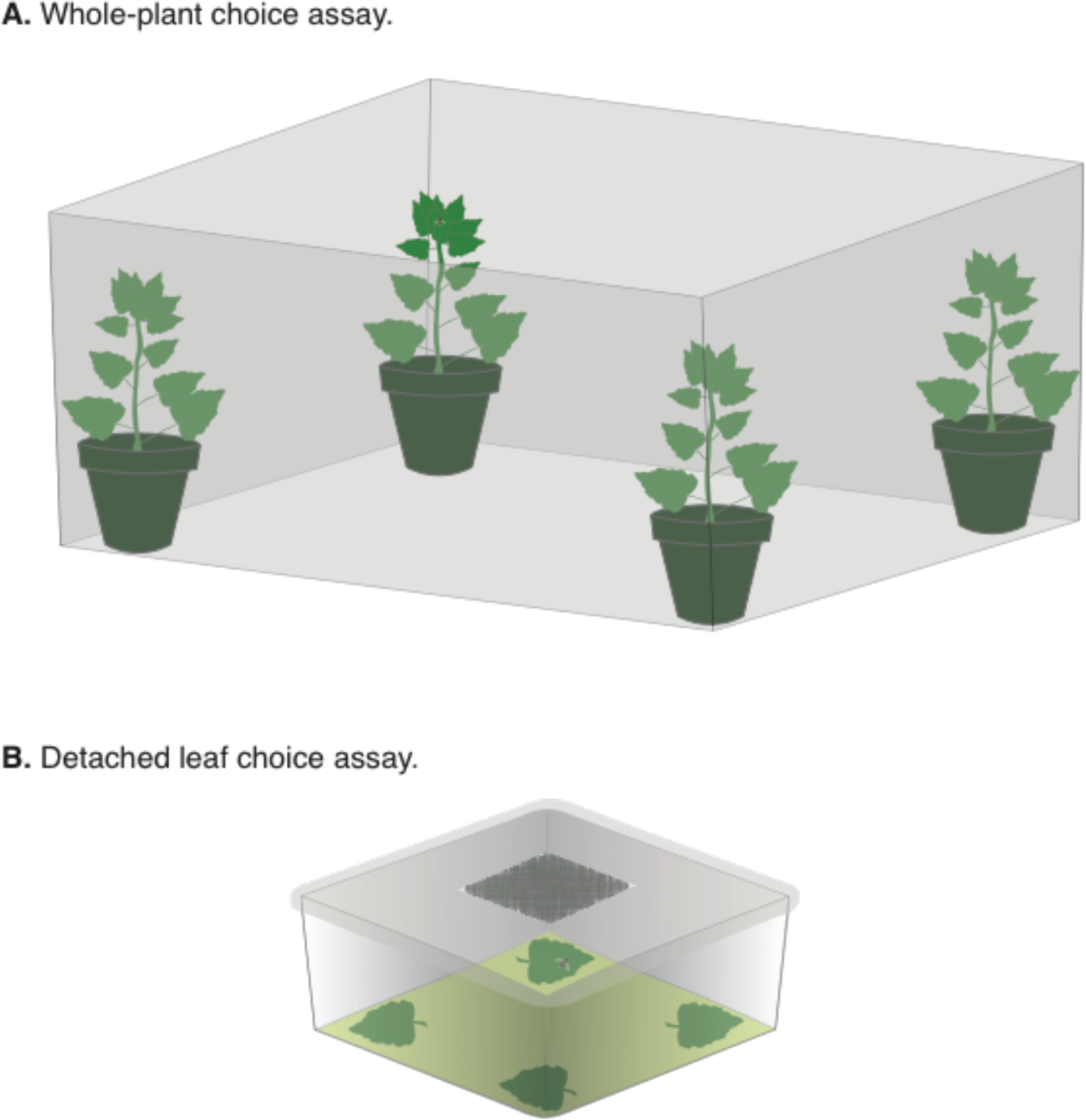
Schematic of experimental design for Host choice experiment I: Sun versus shade-derived bittercress. (**A**) Whole-plant assay depicted (eight replicate trials were conducted; see Materials & Methods, main text). (**B**) Detached leaf assay (fifteen replicate trials were conducted; see Materials & Methods, main text).

**Fig. S2.**
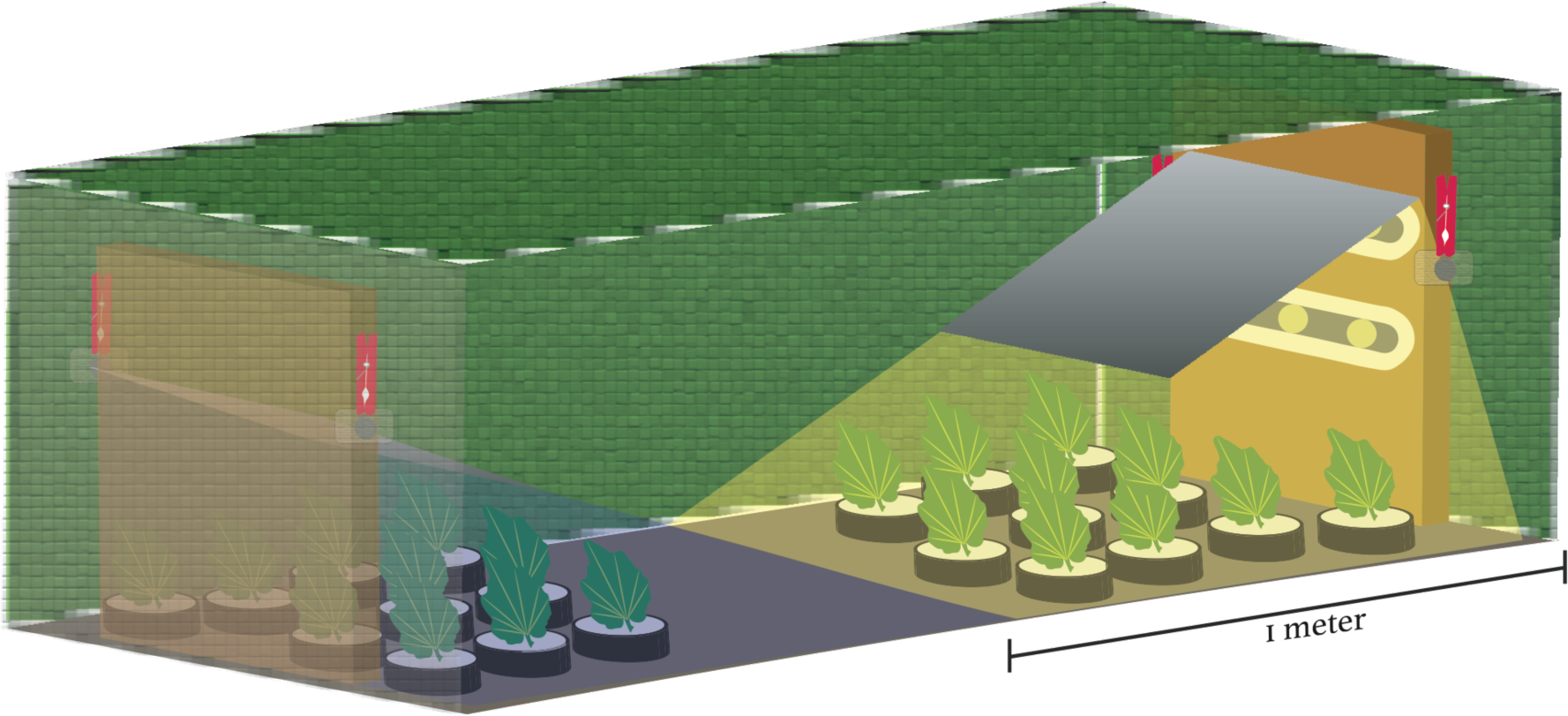
Schematic of experimental design for Host choice experiment II: Effects of light and temperature.

## APPENDIX 3: STATISTICAL SUPPLEMENT

### ALEXANDRE, HUMPHREY ET AL

#### 1. Herbivory survey

##### 1.1 Biological motivation

Herbivore damage (in this case, counts of feeding punctures made by adult females [‘stipples’], leaf mines made by larvae, and eggs laid by adult females) is typically recorded as positive integer counts, and such count data typically exhibit under-dispersion (i.e. ‘zero-inflation’), over-dispersion (‘excess variance’), or both, with respect to expectations of Poisson or Poisson–Gamma mixture (i.e. negative binomial, ‘NB’, models). Our understanding of the foraging ecology of *Scaptomyza nigrita* leads us to expect that dispersion of both of these types is likely. Female flies are choosy, often visiting several leaves before making feeding punctures; females may also avoid leaves entirely, either actively or due to stochasticity in the host sampling process, or to differences in local abundances of foraging *S. nigrita*. All three processes can affect the prevalence of damage among leaves and lead to under-dispersion. The extent to which this is observed may well vary among sites or habitat types.

In addition, once a host has been preliminarily accepted, we expect that variation in the intensity of feeding damage arises from factors perceived subsequent to the initiation of damage. We thus assume that separate (but potentially related) biological processes govern the host acceptance versus the host damage stages of herbivory as measured in our study, and this also may vary across plants, site or habitat type. Below, Figure S1 shows the distribution of stipple and mine counts from our herbivory survey, broken down by habitat type, and from these raw data plots we can already appreciate the strong difference between habitat types in the prevalence (0 versus > 0 counts) as well as the average intensity of damage (distribution of counts > 0).

**Figure S1.**
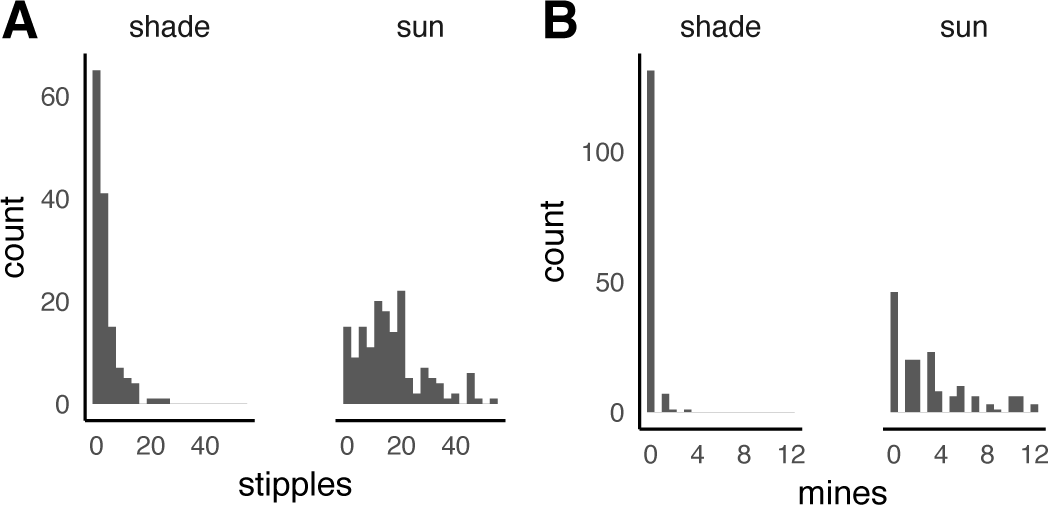
Distribution of (**A**) stippling and (**B**) leaf miner damage on sun and shade-grown bittercress across *n =* 15 sites in a field herbivory survey.

When we compute the ratio of the variance to the mean for stipple and leaf mine counts for each site (Figure S2), we observe variation both within and between habitat types. When examining these two figures, we might suspect that a standard Poisson model may be inappropriate for accounting for the range of variation observed in these data, given that we expect the sample variance to sample mean ratio to be∽ 1 under a simple Poisson process. Instead, we observe (1) ratios > 1 and distributions of ratios that differ between site types (Figure S2; i.e. heteroscedasticity). These two patterns indicate that our count data may *not* be suitably modeled without accounting for zero-inflation and over-dispersion simultaneously. Below we describe our modeling approach using a flexible mixed modeling framework for handling dispersion of both types.

**FIGURE S2.**
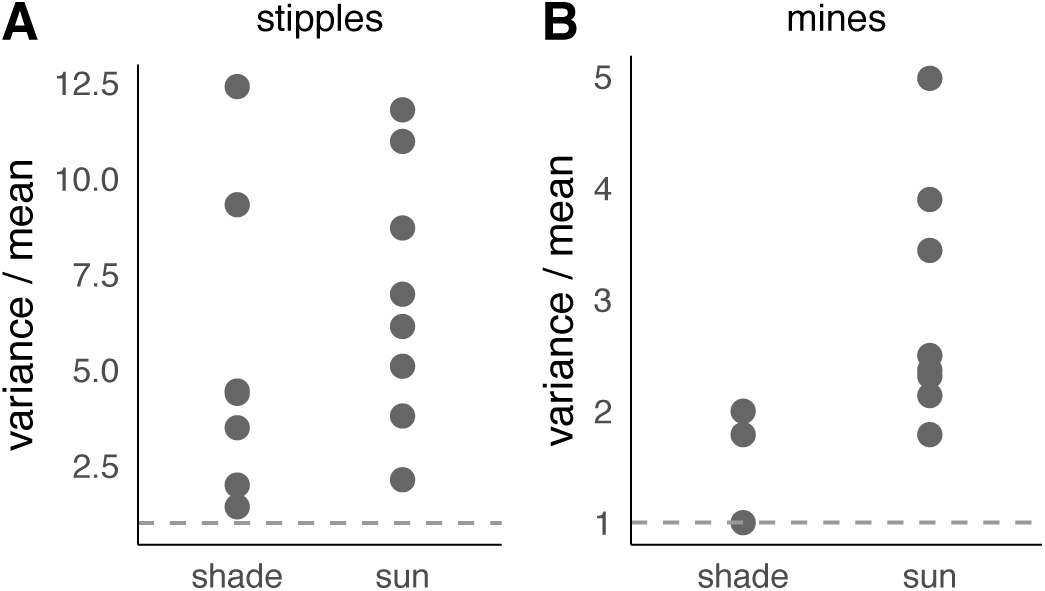
Variance/mean ratios for (**A**) stippling and (**B**) leaf miner damage on sun and shade-grown bittercress.

##### 1.2. Modeling zero-inflation

Zero-inflated models are mixture models composed of a binomial component that models the zero-inflation, and a count component (typically Poisson or NB) that is jointly fit. In such models, the extra zero count compartment results from an ‘artificial’ inflation of the probability density at γ *=* 0 given some unmeasured latent variable that governs the exposure of sampling units to the processes that generate ‘true’ counts [6]. When combined with the probability density function of the Poisson distribution, a zero-inflated Poisson becomes

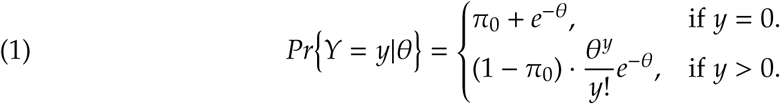

where *θ* is the Poisson mean and *y* is an observed count. The latent variable (*л*_0_) can then be estimated in a binomial model that predicts the compartment from which a given zero count is drawn. We might wish to describe the influence of habitat type on *л*_0_, and to do so we construct a binomial linear model which estimates the zero-inflation coefficient *л*0for each habitat type *k*, using the canonical log it link function:

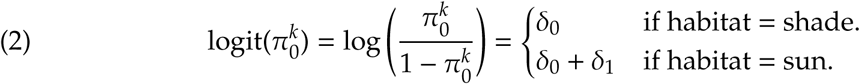

where δ_0_ is the estimated coefficient for the shade habitat, taken as the reference level (i.e. the ‘Constant’ or intercept), to which the effect of the sun habitat (δ_1_) is added. Maximum likelihood coefficient estimates were generated using **R** [4] package gl mm TMB [2, 5, 3]. In the main text Table 1 we report the estimates of *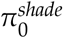* and *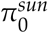* with their 95% (Wald type) confidence intervals on the linear (i.e. the probability) scale instead of the δk estimates on the log it scale.

##### 1.3. Modeling the damage intensity process

Our damage intensity counts are empirically over-dispersed compared to expectations of a Poisson process (Figure S2). This arises when the variance grows faster than the mean, yielding variance to mean ratios > 1. To accommodate this fact, we model counts of herbivore damage (stipples and leaf mines) as a Poisson random variable (*Y*_*i*_) where the Poisson mean (Θ_*i*_) is itself a random variable, which gives rise to the following expression for the conditional probability of *Y*_*i*_ given Θ_*i*_, in the context of zero-inflation:

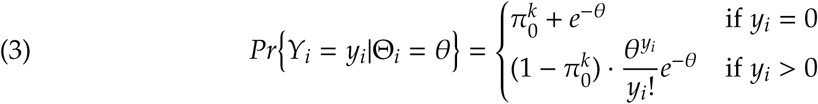

Conceptually, this means that the variance of our random variable γ includes the Poisson variance associated with each Poisson mean Θ_*i*_ as well as additional variance in the distribution of *θ* itself. We adopt the conventional probability density function for *θ* as a Gamma distribution with scale parameter α and rate parameter β [6]:

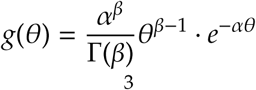

We recover the the unconditional probability of γ _*i*_ by integrating out the *θ* (i.e. averaging over the distribution of Poisson means) to recover the parameterization of the negative binomial (NB) that we will use [6]:

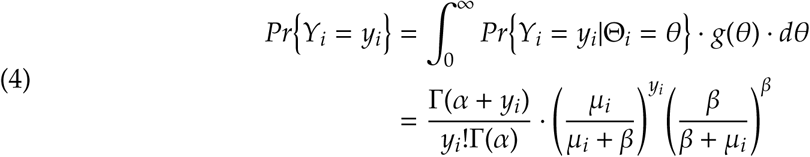

Note that the mean of *g*(*θ*) is *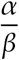* and its variance is*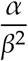*Setting α = β= *ϕ*allows us to re-write the mean as equal to 1 and the variance σ^2^ as *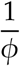*. This gives the expectation of γ _*i*_equal to μ_*i*_ and the variance equal to 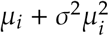. Thus, the ‘dispersion parameter’ of the NB, σ^2^(or alternatively, 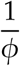), can be interpreted as the variance of the Gamma distribution from which the Poisson means are drawn. When σ^2^→ 0 (and thus*ϕ* → ∞) the negative binomial collapses to the Poisson, where E[γ _*i*_] = Var[γ _*i*_] = μ_*i*_. (Note that the μ_*i*_ of the NB is not equivalent to the *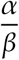*from the Gamma distribution).

The full expression for the unconditional probability of γ _*i*_ becomes (using the *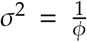* parameterization to be consistent with **R**):

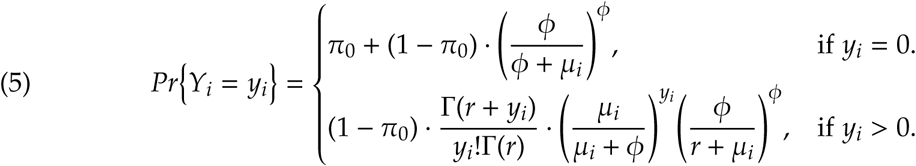

Thus, *Pr {*γ _*i*_} depends on the three parameters, *л*_0_, μ, and Ф In our zero-inflated negative binomial (ZINB) framework, the means of the NB (μ_*i*_) are modeled, via the log link function, as a linear function of four predictor variables:

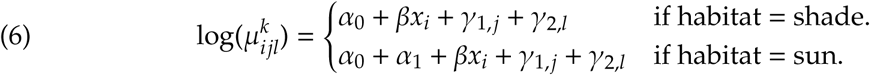

Here, α _1_ indicates the fixed effect of the sun habitat type; γ_1,*j*_ represents a random effect of stem-to-stem variation; and γ_2,*l*_ represents a random effect of site-to-site variation, where *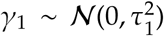* and *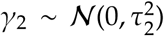* β captures the effect of area (*x*_*i*_) of individual leaves. We sampled two leaves from each of ten plants at all 15 sites, giving a total of *n* = 298observations (two data points were removed due to missing leaf area data). Thus, we have five total parameters to estimate, corresponding to each level of α _*i*_ and the single βterm for the stipple model, as well as the group-level standard deviations for plant stems (τ_1_) and sites (τ_2_).

##### 1.4 Accounting for heteroscedasticity

Differences in the average variance to mean ratio (e.g. Figure S2) across habitat types indicates that certain sub-sets of the data may behave more like a pure Poisson (e.g. Figure S2B, shade) while another subset may behave more like an NB process. Analogous to a Gaussian process, when focal data sub-sets have unequal variances one can estimate separate residual variance terms for each set in order to improve goodness-of-fit and achieve more reliable parameter estimates. In the context of an NB process, accounting for heteroscedasticity can take the form of estimating separate dispersion parameters (*ϕ*) for focal factor levels. In this case, we utilize the flexibility of package glmm TMB to generate maximum likelihood estimates of the NB dispersion parameter *ϕ* as a linear function of habitat type, using a log link:

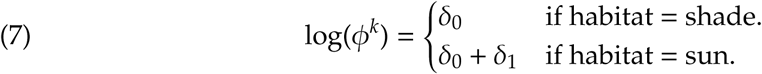

##### 1.5 Model diagnostics and comparisons

Below we illustrate our modeling approach by presenting a justification for using NB over Poisson models, as well as our inclusion of factor-specific zero-inflation and NB dispersion parameters, focusing on the stipple counts from the herbivory survey as an example. We do so by calculating several test statistics that help us judge whether the model construction produces simulated counts which adequately reflect the true but unknown count generating process that we observed in our studies. From each model fitted under maximum likelihood, we generated *n* = 1000 predicted distributions of count data and calculate the proportion of simulations that are more extreme than the observed value. This serves as an empirical *p*-value calculated as *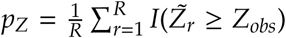*where *I*(.) is the indicator function that equals 1 if the condition is true and 0 otherwise, *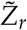*is the test statistic of the simulated data for replicate *r*, and *Z*_*obs*_ is the observed value of the same test statistic. If 𝒫_*Z*_ < 0.05 or > 0.95, we are inclined to think that the model provides an unsatisfactory representation of the underlying data generating process through the lens of a given test statistic. Our test statistics are (1) the proportion of zeros in the data 𝒫 (0), (2) mean(γ), and (3) max(γ). These were chosen as ways to characterize goodness-of-fit the estimated data-generating process at both ends of the data distributions. All raw data and **R** code to fit all models presented in this Appendix can be found in the Dryad data repository (doi pending).

###### 1.5.1 ZI justification

To justify our use of a ZINB model for the herbivory survey data, we first constructed Poisson-only models and compared them to global or habitat-specific ZI-Poisson models. Model checking (Figure S3) reveals that the proportion of zeros under the fitted Poisson-only model simulations is not representative of the observed data(𝒫_*Z*0_ = 0.03 for shade sites, 𝒫_*Z*0_10^−3^ for sun sites). In other words, the model simply fails to accurately represent the prevalence of damage across the dataset. When introducing a global zero-inflation term *л*_0_, model fit greatly improves at the low end, as 𝒫_*Z*0_ increases to-0.85 for sun sites (Figure S4), and the overall goodness-of-fit improves substantially when judged by AICc (ΔAICc = 129; Table S1). But we are still doing a poor job at capturing𝒫 (0) in shade habitats, suggesting that habitat-specific estimates of *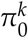* may help. Indeed, modeling habitat-specific ZI improves the goodness-of-fit in shade habitats specifically (𝒫_*Z*0_ ∽ 0.40 for both habitat types; Figure S5). Including an additional model parameter is also statistically favored overall (△AICc = −1.8; Table S1).

**Figure S3.**
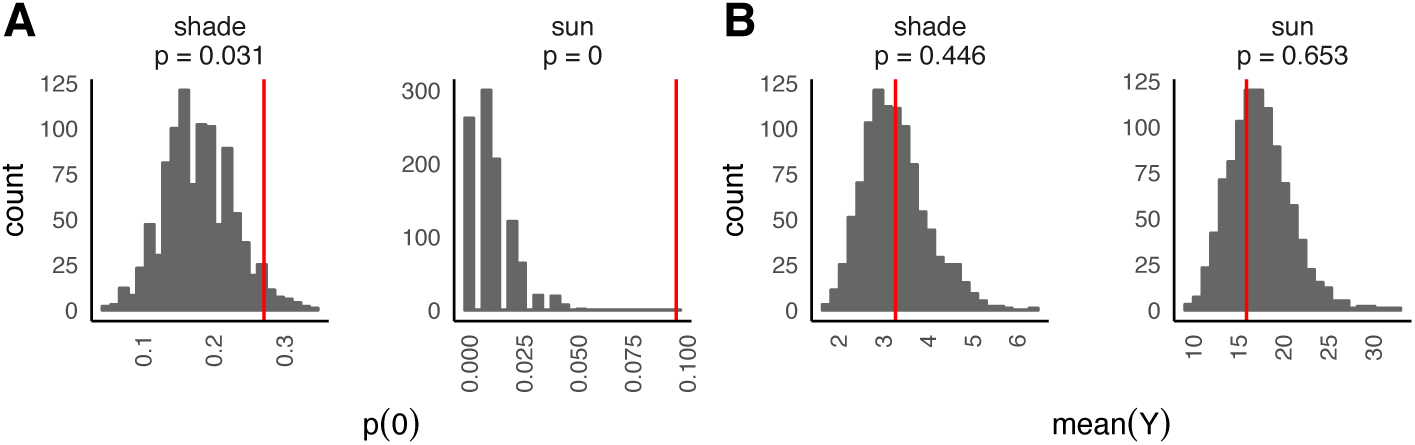
Predictive model checks of (**A**) proportion of zero counts 𝒫 (0) and (**B**) mean(γ) for Poisson-only model of stipple counts. Observed values are indicated by red vertical lines.

**Figure S4.**
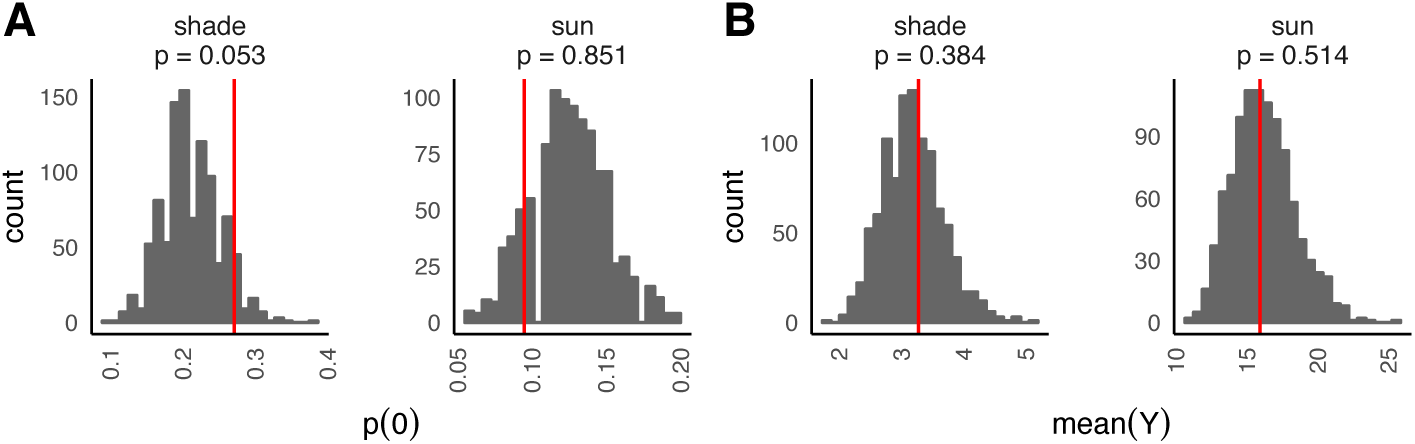
Predictive model checks of (**A**) proportion of zero counts 𝒫 (0) and (**B**) mean(γ) for ZI-Poisson model of stipple counts with single global ZI parameter *л*_0_. Observed values are indicated by red vertical lines.

**Fig. S5.**
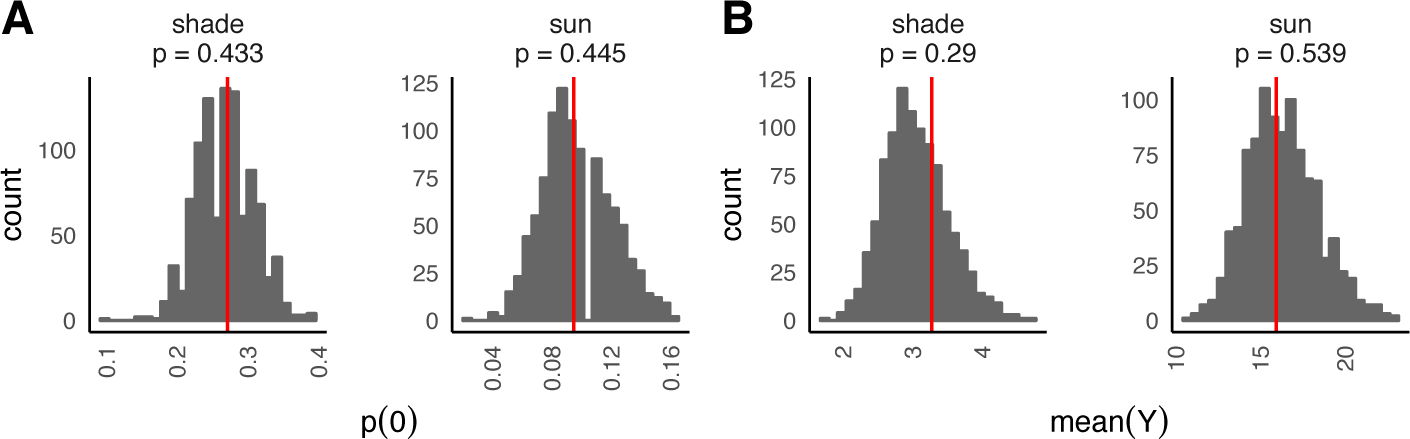
Predictive model checks of (**A**) proportion of zero counts *p* (0) and (**B**) mean(γ) for ZI-Poisson model of stipple counts with habitatspecific ZI parameters *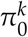*. Observed values are indicated by red vertical lines.

When we conduct a third predictive check to see how the improved model behaves at the high end, via calculating max) (γ), we see that the Poisson model fails to capture the difference between sites in the maximum counts observed (Figure S6).

**Figure S6.**
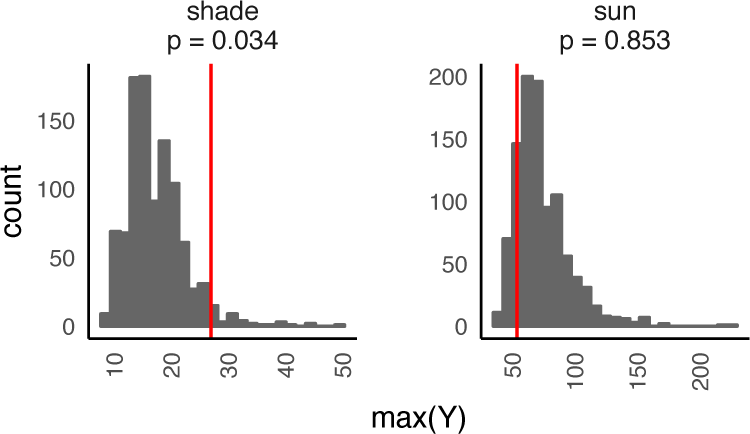
Predicted versus observed max(γ) for ZI-Poisson model of stipple counts with habitat-specific ZI parameters *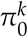*. Observed values areindicated by red vertical lines.

###### 1.5.2 Zero-inflated negative binomial

Model checking in this way shows that Poisson models, even when including habitat-specific ZI terms, can only take us so far in predicting the observed data across its entire range. The lack of fit of the *same data used to estimate the model* indicates lack of flexibility in the model. Because we care about accurately modeling zero counts (i.e. damage prevalence) as well as damage abundance patterns, we explorea set of NB models in order to test whether introducing an additional latent parameter (NB dispersion parameter Ф) improves the ability of the model to capture habitat-specific patterns of herbivory.

Specifically, we compared NB-only to a series of NB models containing a single global ZI, habitat-specific ZI, as well as habitat-specific dispersion parameters with and without consideration of ZI (Table S1). Ultimately, the best model that gives both the lowest AICc (Table S1) as well as the most representative 𝒫_*Z*_ estimates is a NB model with a global*л*_0_ and habitat specific Ф^*k*^. Specifically, this model performs at least as well as the twolevel ZI-Poisson while also fitting better at the upper range, as indicated by the more representative range of max(γ) (Figure S7).

This same analysis procedure was applied for the leaf miner data from the herbivory survey. These results are not displayed here but can be found in the Dryad repository (doi pending).

**Table S1.** Goodness-of-fit statistics for Poisson and NB models with and without accounting for ZI and dispersion.

| family | $\Delta\text{AICc}$ | df | $\pi_0^*$ | NB $\phi^*$ | $p(0)^{\text{shade}}$ | $p(0)^{\text{sun}}$ | $\text{mean}(Y)^{\text{shade}}$ | $\text{mean}(Y)^{\text{sun}}$ | $\text{max}(Y)^{\text{shade}}$ | $\text{max}(Y)^{\text{sun}}$ |
| --- | --- | --- | --- | --- | --- | --- | --- | --- | --- | --- |
| NB | 0 | 7 | 1 | 2 | 0.59 | 0.38 | 0.41 | 0.48 | 0.31 | 0.78 |
| NB | 1.8 | 8 | 2 | 2 | 0.47 | 0.43 | 0.44 | 0.47 | 0.36 | 0.77 |
| NB | 19.3 | 7 | 2 | 1 | 0.44 | 0.45 | 0.34 | 0.56 | 0.09 | 0.93 |
| NB | 20.6 | 6 | 1 | 1 | 0.09 | 0.80 | 0.39 | 0.51 | 0.08 | 0.94 |
| NB | 63.3 | 5 | 0 | 1 | 0.07 | < 0.001 | 0.46 | 0.62 | 0.32 | 0.99 |
| Poisson | 72.5 | 6 | 2 | – | 0.43 | 0.45 | 0.29 | 0.54 | 0.03 | 0.85 |
| Poisson | 74.3 | 5 | 1 | – | 0.05 | 0.85 | 0.38 | 0.51 | 0.05 | 0.90 |
| Poisson | 204.0 | 4 | 0 | – | 0.03 | < 0.001 | 0.45 | 0.65 | 0.20 | 0.94 |

\* 1 = single global parameter; 2 = habitat-specific parameters; 0 = not estimated; – = not applicable

##### 2 Herbivore choice tests I: Sun versus shade derived bittercress

###### 2.1. Models for choice experiments

Using NB generalized linear mixed models (GLMMs), we model variation in stipple and egg counts on plants arising from source habitat (sun versus shade) and leaf attributes (as fixed effects), as well as ‘structural’ random effects of plant ID nested within cage ID (i.e. replicate) to capture experimental design constraints that determine the level of independence among data points. We do not consider Poisson models in all subsequent analyses on the basis of far worse model fits observed for stipple and leaf mine data, above. A priori, we disfavored considering ZI models because we assumed that *S. nigrita* females had suicient time to sample all available plants, and that variation in the intensity (rather than prevalence) of herbivory would drive most of our patterns. However, in all cases we considered whether including source habitat-specific estimates of *ϕ*^*k*^ empirically improved model fit via AICc (see Results and Table 1 of main text).

Our mixed model for stipple and eggs counts, for both the whole-plant and detached leaf assays, takes the following form:

**Figure S7.**
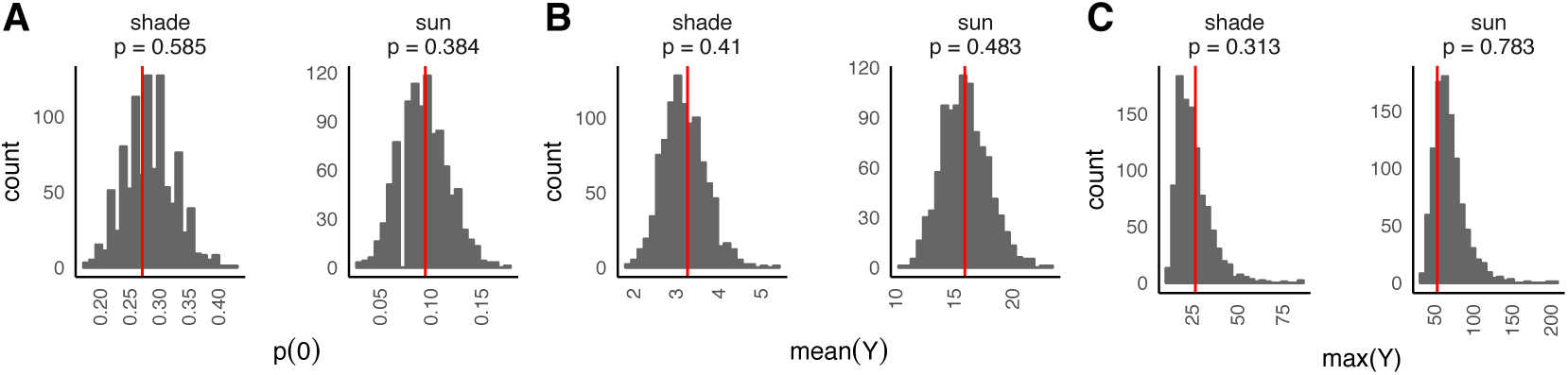
Predictive model checks for best-fitting ZINB model of stipplecounts, with habitat-specific dispersion parameters *ϕ*k and global ZIparameter *л*0. Observed values are indicated by red vertical lines.

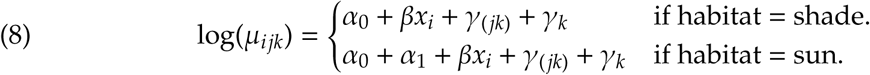

In this model, μ_*ijk*_ is the estimate for each leaf *i*, α_0_ and α are as in Eqn. 6, and f3 is leaf position along stem (low to high) for the whole-plant model or leaf area (mm^2^) for the detached leaf assay; finally, γ_*k*_ represents a random effect of cage ID (*k*) and γ _(*jk*)_ representsa random effect for plant ID *j* nested within each level of cage ID *k*; 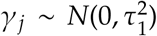 and 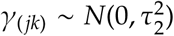 A walk-through of model comparisons and goodness-of-fit comparisonsy for this assay can be found in associated **R** notebooks (Dryad data repository).

##### 3. Herbivore choice tests II: Effects of light and temperature

The NB model structures for this set of choice experiments were similar to those described in above except for an expanded random effects structure. As for the previous herbivore choice models, all models described below are NB-only GLMMs. The choice trials in 2014 and 2015 were conducted slightly differently, which means that each year’s data calls for slightly different random effects. We present the analyses of each dataset separately, then jointly.

###### 3.1 2014 Trials

The 2014 field and lab trials were conducted in two temperature environments simultaneously, with one cage held in each. This gives a structure of temperatureenvironment (γ _(*lk*)_, *n* = 2) nested within trial (γ _*l*_, *n* = 6 each for the 2014 field and lab assays). Nested within temperature environment is cage, but since we have only a single level of cage for each temperature environment per trial, this level is irrelevant for the 2014 dataset. However, we include side-of-cage (γ _(*kj*)_, *n* = 2; left or right, arbitrarily) to control for pseudo-replication at the level of the main treatment effect (i.e. Light environment, light versus dark), which was applied with randomization to the sides of each cage. Each random effect is 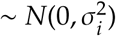. The full model is given below using the above notation for the random effects. The number of independent data points in the 2014 field and lab trials is 24 each (2 sides per cage, 1 cage per trial, 2 temperature settings per trial, and 6 trials).

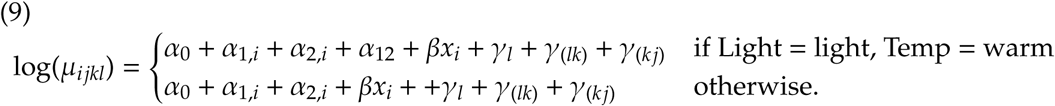

We set the first levels of coefficients for the fixed effects of Light (α _1,1_ = dark) and Temperature (α _2,1_ = cool) equal to zero since these two states are incorporated into the Constant (α _0_, i.e. the reference level of the model). Our experiment was designed to allow us to fit an interaction term α _12_ which estimates how much the effect of Light is impacted by the Temperature environment (thus α _12_ has a single level). We fit models with the same structure for field and laboratory trials, and we consider condition-specific *ϕ* estimates when conducting model comparisons. Specifically, we estimated whether *ϕ* specific tolight condition and/or temperature condition improved model fit by accounting for differential distributions of overdispersion analogous to Eq. 7 (with an additive (on the log scale) impact of temperature treatment).

###### 3.2 2015 Trials

In the 2015 trials, we used bittercress leaves collected from both sun and shade habitats, and our randomization scheme ensured equal representation of sun- and shade-derived leaves across all treatments. We did this to test whether *S. nigrita* would exhibit preference for shade-derived bittercress in the context of our temperature and light manipulations. In our analysis, we included leaf source (sun v. shade) as an additional fixed factor. Our model structure was thus

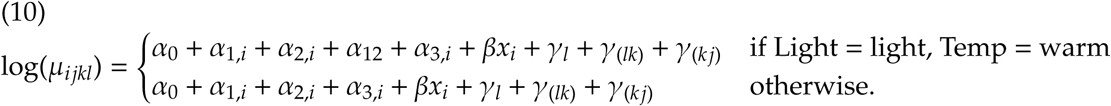

All coefficients are as in the 2014 trials above, except that the term α _3,*i*_ is for source habitat, and the first level (α _3,1_ = shade) was set to 0 and was thus incorporated into the intercept term α _0_. Thus, this model estimates only a single coefficient for the effect of habitat.

The experimental procedures also differed slightly in 2015. We placed two cages in one environmental chamber which was set to a single temperature per trial, and individual trials were conducted at different temperatures sequentially. This implies a structure ofcage (γ _*kl*_, *n* = 2) nested within trial (γ _*l*_, *n* = 5), along with the side-of-cage nested within cage (γ _(*kj*)_, *n* = 2). The number of independent data points in the 2015 trials is thus 40(2 sides per cage, 2 cages per room, 10 trials [5 in each temperature regime]). While the model structure is the same as in Eqn. 8, the meaning of the random effects (and their coding in the design matrix) are slightly different. Below we discuss how we reconciled the random effects to analyze both years’ data together.

###### 3.3. Combined 2014-2015 Analysis

We reconcile the slight distinctions between 2014 and 2015 datasets by including trial-year as a composite random effect, now re-coded to reflect each experiment conducted in a given room (compared to a trial containing two separate temperature settings, as in the 2014-only analysis). Nested within the new trial factor is cage, which has n = 1 for 2014 and n = 2 for 2015; side-of-cage is modeled in the same way as above. This model structure is now identical to Eqn. 10.

For the combined analysis, we drop the term for plant source habitat (α _3_) since the 2014 trials were not designed to examine plant source habtat. Additionally, we model botha continuous and discrete versions of the Temperature factor: the discrete model is the same as Eqn. 8, while in the continuous form, α _2,*j*_ is replaced with β _2_ γ _*j*_, where γ _*j*_ is the temperature measured at the level of cage for each trial separately (Appendix B, Fig. 2C). Additionally, the fixed interaction term α _12_ is now replaced with an interaction modeled by the expression β γ _*j*_(α _1,*j*_), which captures how the effect of light (α _1,*j*_) changes as a function of cage temp (γ _*i*_); since α _1,*i*_ has one level (the first level is set to 0), estimating this interaction termβ _3_ adds only a single parameter to the model. The full model is thus:

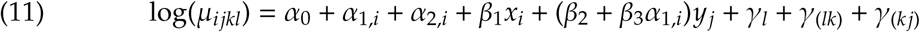

Because the full model with the continuous temperature interaction term (β_3_) exhibited such a poor fit to the data as judged by AIC (see below), we report the coefficient estimates for the nested model containing only the additive effect of the continuous temparture term (β_2_; Table S2).

**Table S2.**
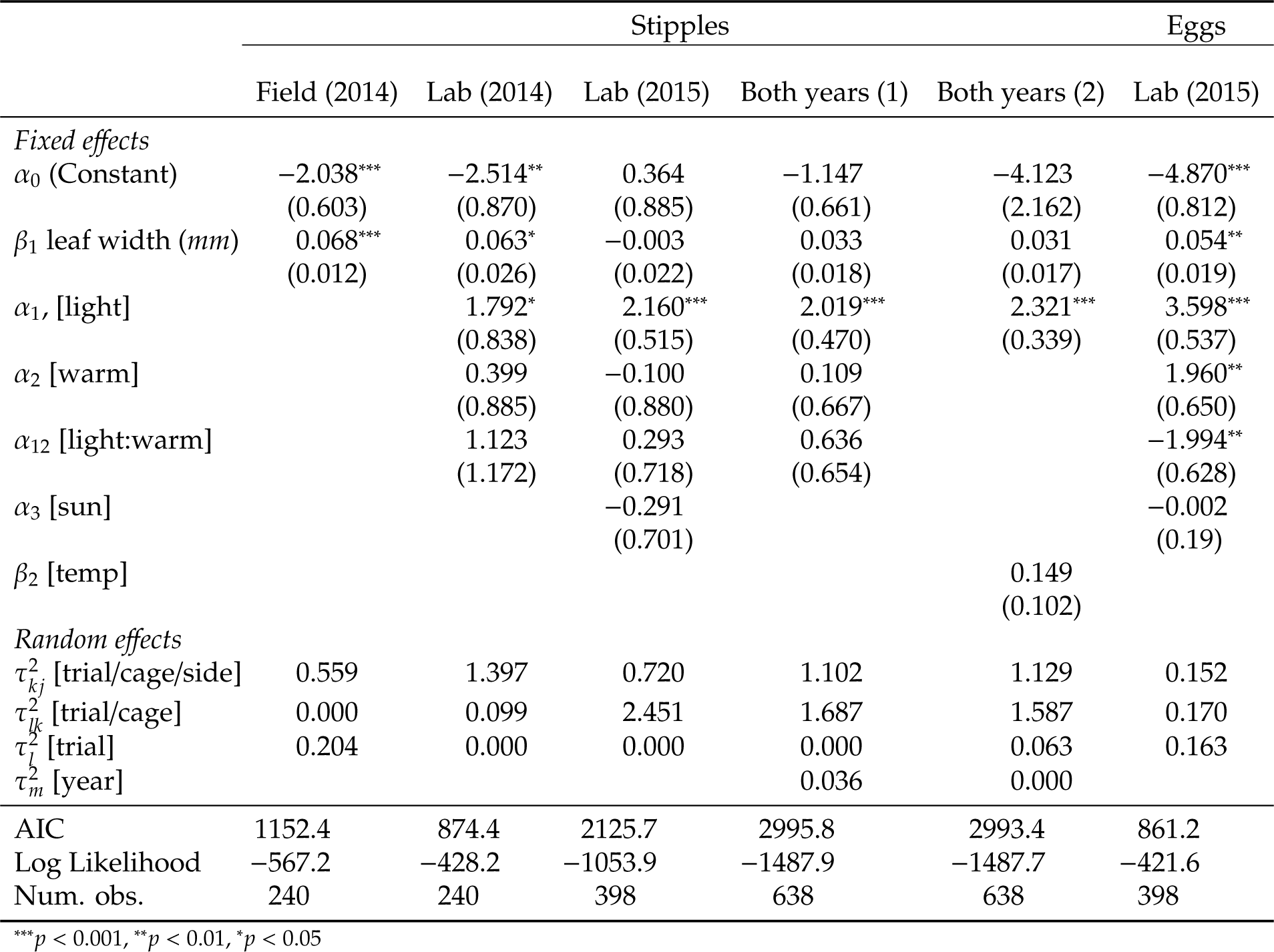
Coefficient estimates for all light–temp choice experiments.

## Appendix 4

### Temperature profiles for field and laboratory habitat preference choice tests

**Fig. S1.**
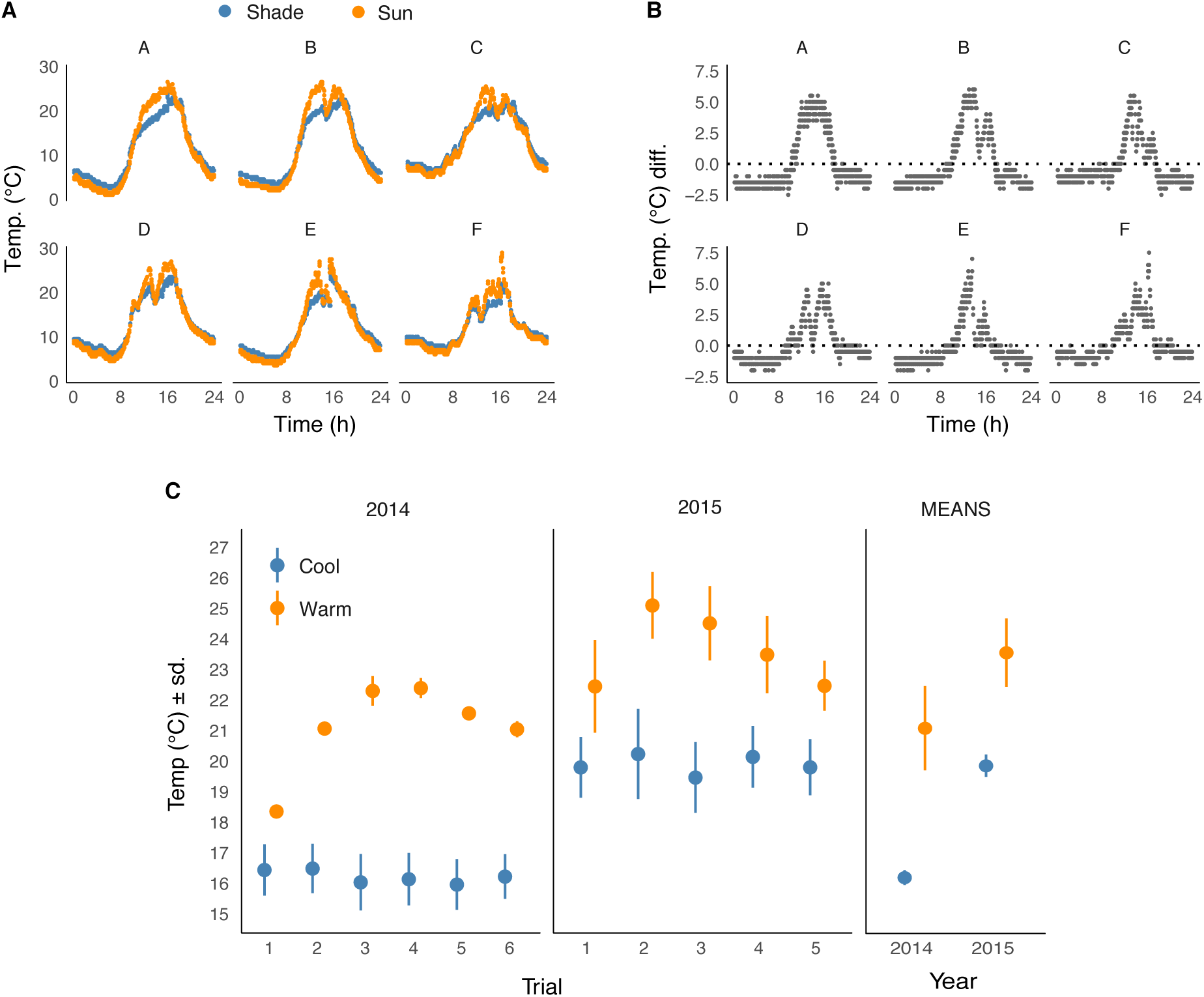
Temperature profiles for field and laboratory habitat preference choice tests. (**A)** Temperature profiles for field cages (from 2014). Both ‘Sun’ and ‘Shade’ cages were equally masked from natural sunlight but were either sun-exposed or canopy shaded in order to confer different temperature profiles. We collected a full 24 h of temperature data during each trial (ordered in time and labeled A–F), which took place for 24 h beginning at 1100 h. 0 h represents midnight. (**B**) Differences in temperature between sun-exposed and canopy-shaded assay cages (sun – shade), showing a maximal difference of 5 °C in mid-afternoon during each trial. (**C**) Average temperature for each laboratory trial for 2014 and 2015 (left; ordered in time and labeled 1–6), and the mean (right; ± 1 standard deviation) over all trials for each year. 2014 trials used two environmental chambers, while 2015 trials alternated between warm and cool trials at two-day intervals (see Methods in the main text for details).

## Appendix 5

### Distribution of stipples and eggs on sun- and shade-source plants in the abiotic habitat choice trials

**Fig. S1.**
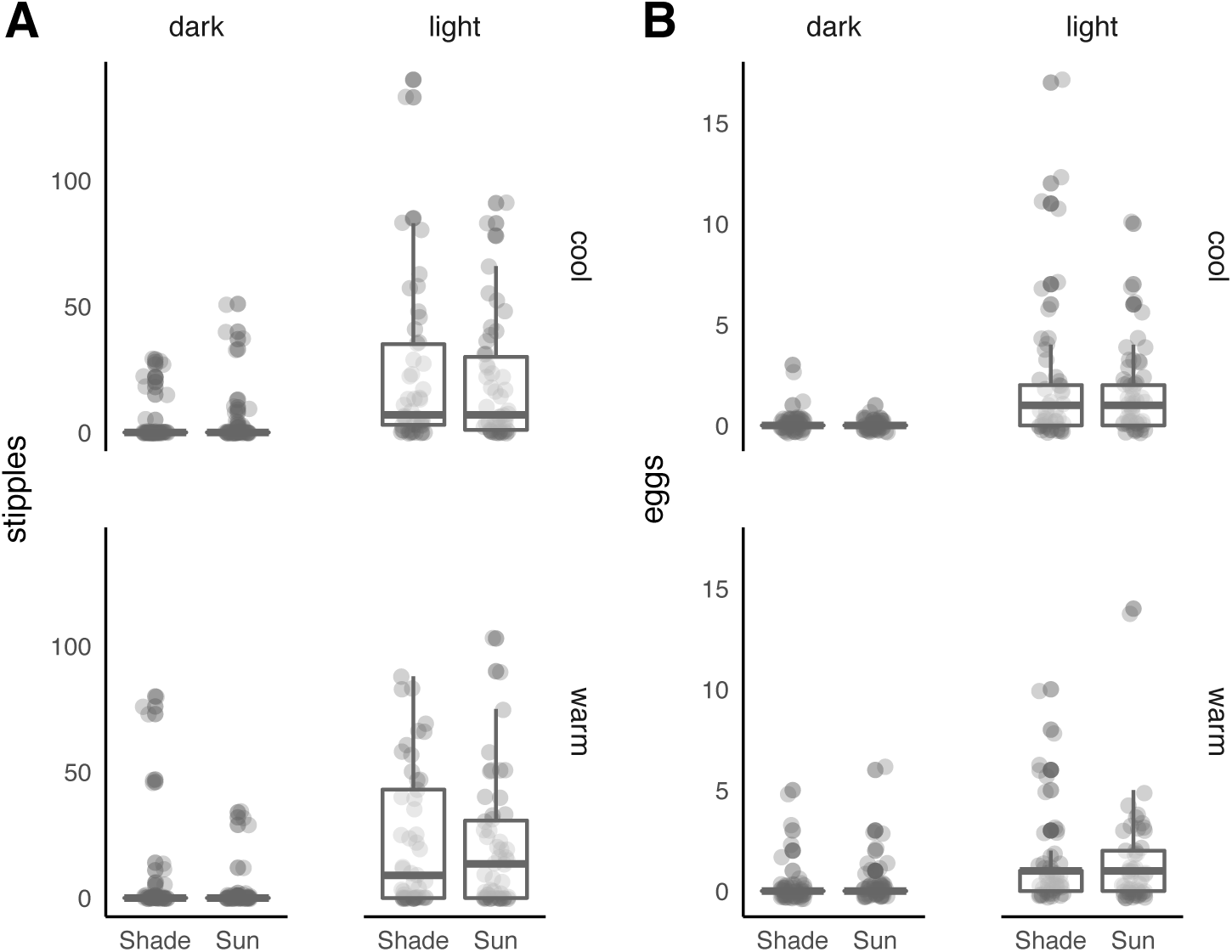
Sun and shade-source plants received indistinguishable amounts of stipples (**A**) and eggs(**B**) across light and temperature treatments in the abiotic habitat choice trial with adult female *S. nigrita* flies.

Author contributions
NMA, ADG, JL, HAA, and NKW designed the experiments; NMA, ADG, JL, HAA, and JF collected the data; PTH and NMA analyzed the data; NMA, PTH and NKW wrote the paper. All authors contributed to editing the submission

